# Delta/Notch signaling level is decoded through transcriptional burst duration during muscle progenitor specification

**DOI:** 10.64898/2026.10.30.763040

**Authors:** Emma Leroux, Olivier Josse, Savannah Moinet, Cyrille Surbled, Aurélien Guillou, Mounia Lagha, Hadi Boukhatmi

## Abstract

The ability of progenitor cells to interpret different levels of signaling pathway activity is fundamental to the generation of cellular diversity during development. Muscle progenitors, for example, are exposed to different levels of signals from their microenvironment. However, how they decode these quantitative differences to regulate their cellular behaviors during myogenesis remains poorly understood. Here, we addressed this question during the specification of two populations of *Drosophila* adult muscle progenitors (AMPs). We found that, despite being spatially adjacent, indirect flight muscle (IFM) and direct flight muscle (DFM) progenitors receive distinct levels of Notch signaling. We further show that the spatial distribution of epithelial Delta contributes to differential Notch activation. Higher Delta levels adjacent to IFM progenitors promote stronger Notch signaling. To investigate how these quantitative differences are translated into transcriptional responses, we developed a Notch-responsive MS2 reporter to monitor transcription in living AMPs. Our measurements reveal that Notch-dependent transcription occurs in stochastic bursts, with IFM progenitors displaying longer but less frequent bursts than DFM progenitors. Strikingly, equalizing epithelial Delta levels abolished differences between the two populations in both Notch-dependent transcriptional activity and the protein levels of Notch target genes. Together, our findings suggest that quantitative differences in signaling may be translated into distinct transcriptional outputs through modulation of transcriptional bursting. This may represent a mechanism by which a single signaling pathway can be deployed to regulate diverse cellular responses during development.

## Introduction

The establishment of cellular diversity during development relies on the ability of progenitor cells to perceive and interpret extracellular signaling cues and execute distinct gene expression programs. Because only a limited number of signaling pathways govern most developmental processes, the same pathways are repeatedly used to regulate diverse cell fate decisions [1]. Increasing evidence indicates that signaling dynamics and the level of pathway activation provide an additional layer of regulation. These features enable the same signaling pathway to convey distinct regulatory information that directs different cell fate decisions [2]. However, how these signaling features are decoded into distinct transcriptional patterns that ultimately specify different cell fates remain largely unknown.

Notch signaling regulates numerous developmental processes, including progenitor maintenance, proliferation, differentiation, and its disruption is associated with a wide range of developmental disorders and diseases [3]. Notch signaling is initiated through direct cell-cell interactions, whereby transmembrane ligands of the Delta-Serrate-Lag (DSL) family bind the Notch receptor and trigger two successive proteolytic cleavages that release the Notch intracellular domain (NICD). NICD subsequently translocates to the nucleus, where it associates with the DNA-binding factor CSL (CBF1/Suppressor of Hairless/Lag-1) to activate transcription of Notch target genes. Unlike many developmental signaling pathways, Notch signaling is remarkably linear and lacks an intermediate amplification step. Consequently, the amount of NICD generated is expected to scale directly with receptor activation. Thus, differences in ligand availability are predicted to produce proportional differences in the amount of NICD reaching the nucleus and, consequently, in Notch signaling output [4].

Initially, Notch signaling was viewed as a binary pathway in which receptor activation induces the expression of specific target genes that, in turn, drive cell fate specification. However, recent studies have challenged this classical view by showing that the dynamics of Notch signaling can generate distinct transcriptional responses and developmental outcomes. For example, oscillatory expression of Notch target genes promotes neural stem cell self-renewal, whereas sustained expression drives neuronal differentiation [5]. Furthermore, oscillatory expression of the Notch ligand Delta-like 1 has been shown to be essential for maintaining the balance between self-renewal and differentiation of mammalian muscle stem cells [6]. Consistent with this concept, Dll1 and Dll4 activate Notch through distinct temporal modes, with Dll1 eliciting pulsatile Notch activation and Dll4 inducing sustained signaling. These different signaling dynamics activate distinct transcriptional programs and ultimately promote opposite myogenic cell fate outcomes [7]. Together, these studies highlight that, beyond a simple ON/OFF response, differences in Notch ligand availability and signaling dynamics contribute to distinct cell fate decisions.

The development of quantitative methods for monitoring transcription, such as single-molecule fluorescence in situ hybridization (smFISH) and MS2-MCP live imaging, has revealed that genes are not transcribed continuously [8]. Instead, transcription occurs in discrete stochastic bursts of activity separated by periods of transcriptional inactivity [9]. An increasing number of studies have shown that transcriptional responses to Notch signaling also occur in stochastic bursts [10–12], raising the possibility that modulation of these transcriptional dynamics could contribute to establishing distinct levels of target gene expression and, consequently, distinct cellular responses to Notch signaling.

The diversity of the *Drosophila* musculature provides an excellent model for investigating how quantitative differences in signaling pathway activity contribute to progenitors specification during development. Adult flight muscles comprise two morphologically and functionally distinct muscle types: the fibrillar indirect flight muscles (IFMs) and the tubular direct flight muscles (DFMs). Both muscle types arise from a common pool of adult muscle progenitors (AMPs) associated with the wing imaginal disc [13, 14]. During their specification, the wing disc epithelium acts as a transient signaling niche, providing multiple cues that regulate AMP maintenance, proliferation, and differentiation [15, 16]. Among these cues, Notch signaling, activated by the epithelial ligands Delta and Serrate, is essential for maintaining AMPs in an undifferentiated and proliferative state [17–20]. Interestingly, recent single-cell RNA-sequencing analyses revealed that several Notch target genes are differentially expressed between IFM and DFM progenitors, suggesting distinct transcriptional responses to Notch signaling in these two populations [16].

Here, we investigate whether and how different levels of Notch signaling contribute to the specification of IFM and DFM progenitors in vivo. We show that these two progenitor populations receive distinct levels of Notch signaling and that this difference is driven by the spatial distribution of Delta in the signal-sending epithelium. Using a newly developed MS2-MCP-based reporter, we further show that Notch-dependent transcription occurs in bursts and that higher Notch signaling in IFM progenitors is associated with longer transcriptional burst duration. Strikingly, uniform Delta overexpression throughout the epithelium abolishes differences in both Notch-dependent transcriptional activity and target protein levels between IFM and DFM progenitors. We propose that modulation of transcriptional burst duration provides a mechanism by which progenitor cells translate quantitative differences in Notch signaling into distinct developmental programs in vivo.

## Results

### Delta/Notch signaling is required for normal flight muscle development and function

Notch signaling plays multiple roles in *Drosophila* muscle development [17, 19, 21]. Despite its importance, how Notch loss of function affects the final development and organization of adult flight muscles remains incompletely understood [22, 23]. We investigated further the role of Notch during flight muscle development by specifically depleting Notch in AMPs using the *1151-Gal4* driver to express *UAS-Notch^RNAi^*. This strategy selectively impairs Notch receptor in the AMPs without directly perturbing Notch signaling in the surrounding tissues (Figure S1 A and C-D). Analysis of adult flies revealed that, whereas control flies held their wings flat over the abdomen, *1151-Gal4 > UAS-NotchRNAi* adults consistently displayed a held-out wing phenotype (Figure 1A-A’), commonly associated with defects in flight muscle development or function [24]. To determine whether this abnormal wing posture was accompanied by impaired flight ability, we next assessed flight performance using a standard flight assay (Materials and Methods and [25]). While most control flies reached the top of the flight column, Notch-depleted flies showed a severe impairment in flight performance, with the majority landing in the lower half or at the bottom of the column (Figure 1B-C).

**Figure 1.**
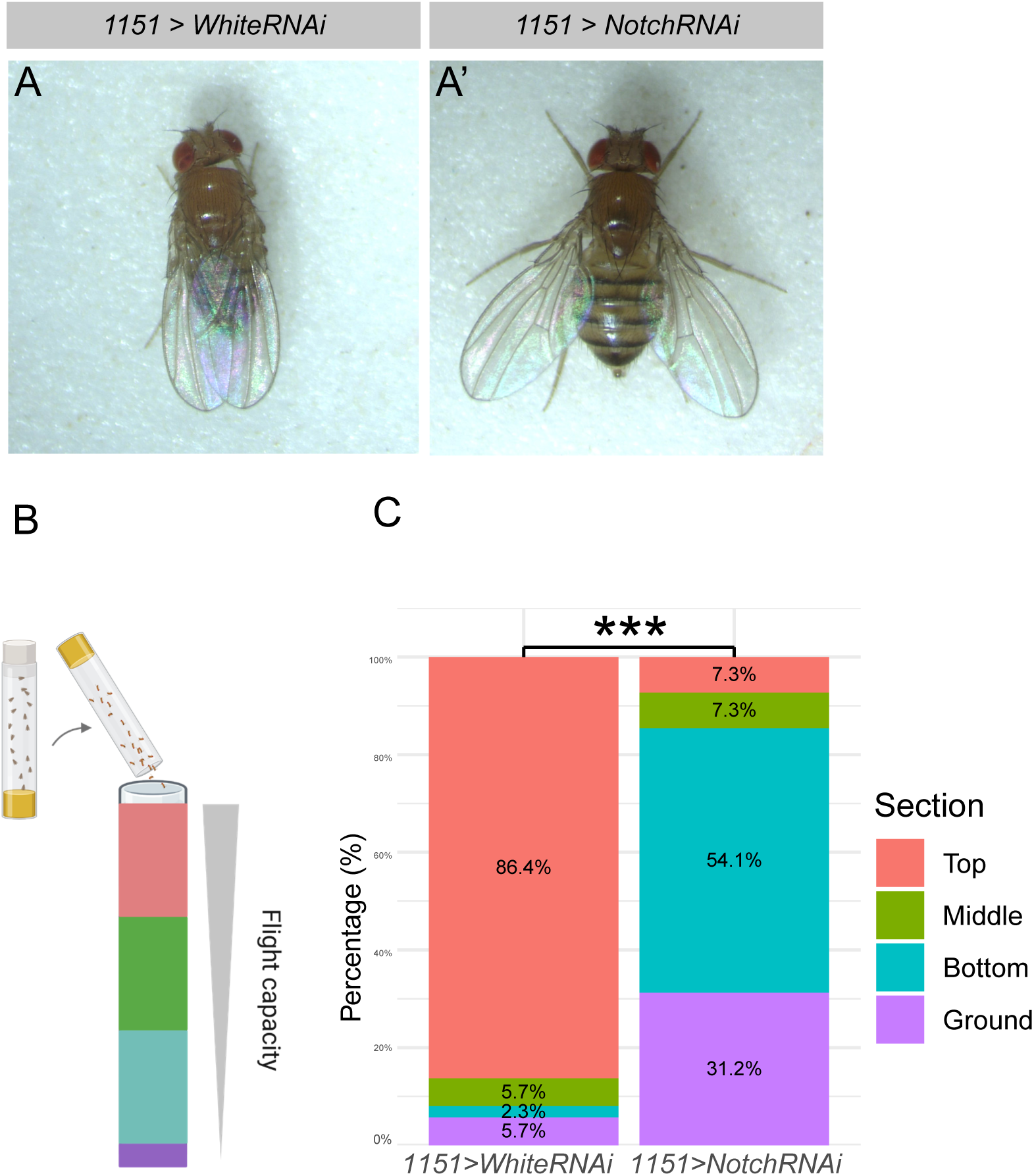
Notch depletion in adult muscle progenitors causes a held-out wing phenotype and impaired flight. (A–A′) Dorsal views of adult flies showing the wing posture of (A) control (*1151 > w^RNAi^*) and (A′) Notch-depleted (*1151 > N^RNAi^*) animals. Notch depletion in AMPs results in a characteristic held-out wing phenotype. **(B)** Schematic representation of the flight assay. **(C)** Quantification of flight performance showing the percentage of flies landing in each section of the flight column (Top, Middle, Bottom, Ground) for control (*1151 > w^RNAi^*) and Notch-depleted (*1151 > N^RNAi^*) flies. Notch depletion significantly impairs flight capacity (***; P < 0.001, χ² test; control *n* = 88 flies; N^RNAi^ *n* = 109 flies; three independent biological replicates).

To determine whether these flight defects were associated with alterations in muscle development, we examined the morphology of the IFMs and DFMs (Figure 2A). Surprisingly, IFMs from *1151-Gal4> UAS-Notch^RNAi^* flies appeared grossly normal and were indistinguishable from those of control animals (Figure 2B-D). Furthermore, we found no significant differences in myonuclear density, the number of IFM fibers, or sarcomere length between control and Notch-depleted muscles (Figure 2C–G and Figure S1 L-N). These results indicate that depletion of Notch in AMPs does not cause overt defects in IFM morphology. Notch signaling in AMPs is activated, at least, by Delta expressed in the adjacent wing disc epithelium [18]. We therefore depleted Delta specifically in the epithelial cells using *Ap-Gal4* (Figure S1 B, E, F and I) and examined the morphology of the adult IFMs (Figure 2G-I). Similar to Notch depletion in AMPs, epithelial Delta depletion did not produce overt structural defects in the IFMs (Figure 2 H-J). Furthermore, myonuclear density and average sarcomere length were not significantly different from those of control muscles (Figure 2L–M).

**Figure 2.**
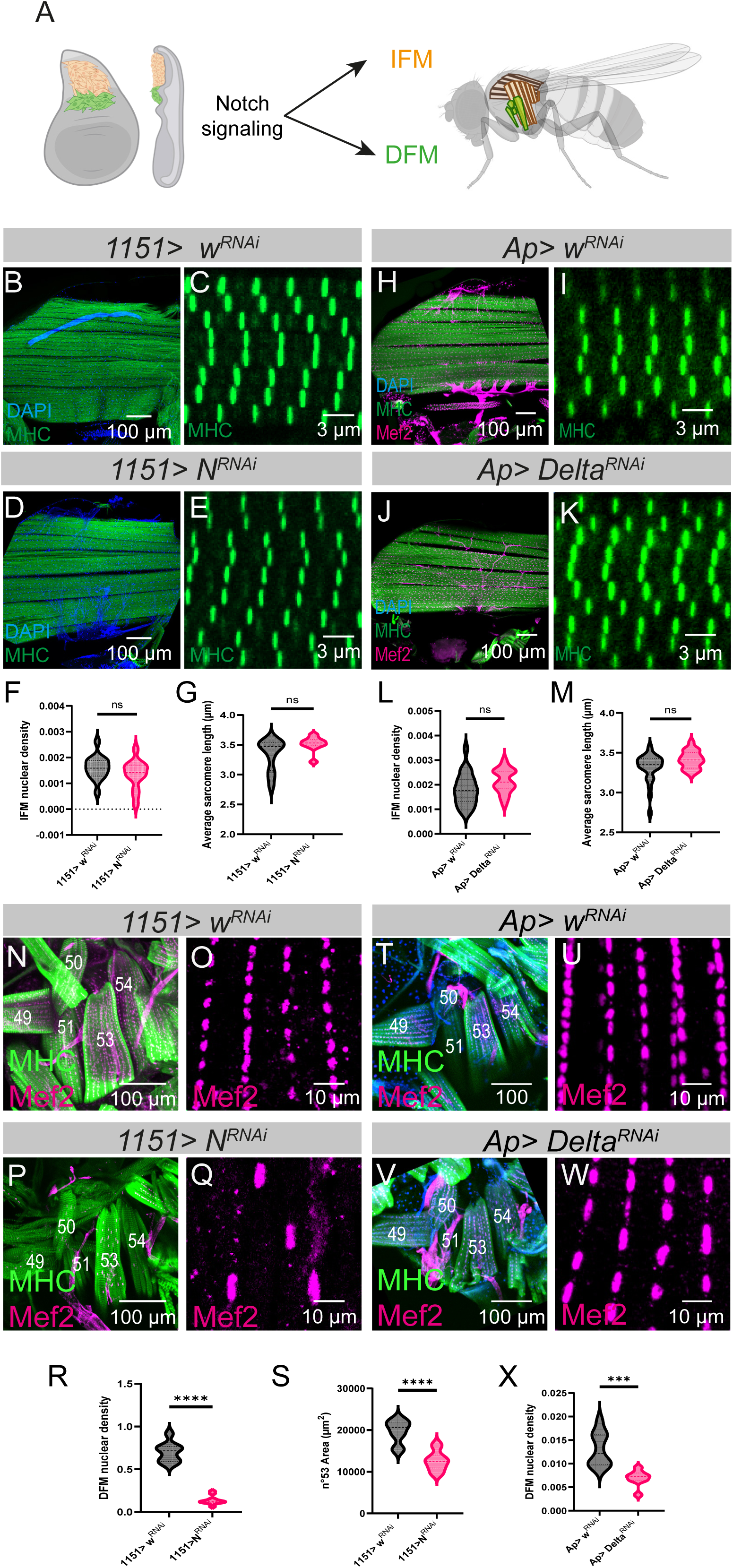
Notch signaling differentially regulates indirect and direct flight muscle development. **(A)** Schematic representation of *Drosophila* flight muscle development. The wing disc adult muscle progenitors (AMPs) are subdivided into two distinct populations that give rise to the adult indirect flight muscles (IFMs, orange) and direct flight muscles (DFMs, green). **(B–E)** Indirect flight muscles (IFMs) stained for Mhc (green) and DAPI (blue) in control (*1151 > w^RNAi^*) (**B, C**) and Notch-depleted (*1151 > N^RNAi^*) (**D, E**) adult flies. Higher-magnification images show Mhc-labelled sarcomeres (**C, E**). **(F)** Quantification of myonuclear density in control and Notch-depleted IFMs revealed no significant difference (ns, *P* = 0.1929, Student’s *t*-test; control, *n* = 15 heminota; N^RNAi^, *n* = 13 heminota; three independent biological replicates). **(G)** Quantification of average sarcomere length showed no significant difference between control and Notch-depleted IFMs (ns, *P* = 0.0720, Mann–Whitney test; control, *n* = 11 heminota; N^RNAi^, *n* = 10 heminota; three independent biological replicates). **(H–K)** IFMs stained for Mhc (green), Mef2 (magenta) and DAPI (blue) in control (*Ap > w^RNAi^*) (**H, I**) and epithelial Delta-depleted (*Ap > Delta^RNAi^*) (**J, K**) adult flies. Higher-magnification images show Mhc-labelled sarcomeres (**I, K**). **(L)** Quantification of myonuclear density in control and epithelial Delta-depleted IFMs revealed no significant difference (ns, *P* = 0.0510, Student’s *t*-test; control, *n* = 24 heminota; Delta^RNAi^, *n* = 19 heminota; three independent biological replicates). **(M)** Quantification of average sarcomere length showed no significant difference between control and epithelial Delta-depleted IFMs (ns, *P* = 0.0701, Mann–Whitney test; control, *n* = 25 heminota; Delta^RNAi^, *n* = 18 heminota; three independent biological replicates). **(N–Q)** Direct flight muscles (DFMs) stained for Mhc (green) and Mef2 (magenta) in control (*1151 > w^RNAi^*) (N, O) and Notch-depleted (*1151 > N^RNAi^*) (P, Q) conditions. Individual DFM fibers are identified by numbers. Higher-magnification images and quantifications were performed on muscle 53 (**O, Q**). **(R)** Myonuclear density in muscle 53 is decreased following Notch depletion (*P* < 0.0001, Student’s *t*-test; control, *n* = 9, preparations; Notch^RNAi^, *n* = 11 preparations; three independent biological replicates). **(S)** Quantification of muscle 53 area shows a significant reduction following Notch depletion (*P* < 0.0001, Student’s *t*-test; control, *n* = 9 preparations; N^RNAi^, *n* = 11 preparations; three independent biological replicates). **(T–W)** DFMs stained for Mhc (green) and Mef2 (magenta) in control (*Ap > w^RNAi^*) (T, U) and epithelial Delta-depleted (*Ap > Delta^RNAi^*) (V, W) conditions. Individual DFM fibers are identified by numbers. Higher-magnification images and quantification were performed on muscle 53 (U, W). **(X)** Quantification of myonuclear density in muscle 53 revealed a significant decrease following epithelial Delta depletion (***, *P* = 0.0004, Student’s *t*-test; control, *n* = 10 preparations; Delta^RNAi^, *n* = 8 preparations; three independent biological replicates).

We next examined the consequences of Notch depletion on DFMs development. In contrast to the IFMs, Notch depletion produced a clear morphological defect in the DFMs (Figure 2 N-P). Although DFM fibers were present, they appeared markedly smaller than those of control flies, with fewer and apparently enlarged myonuclei (Figure 2 O-Q). Consistent with these observations, DFM area and myonuclear density were both significantly reduced following Notch depletion (Figure 2R–S). We then depleted Delta specifically in the wing disc epithelium using *Ap-Gal4* (Figure 2 T-V). Similar to Notch depletion in AMPs, Delta depletion in the signal sending cells produced a marked reduction in DFM myonuclear density (Figure 2 U–W and X). Together, these findings reveal distinct requirements for Delta/Notch signaling during adult flight muscle development. Depletion of Notch or epithelial Delta impairs DFM development, whereas IFMs remain grossly unaffected at the morphological level. Nevertheless, potential defects in IFM contractility or metabolism cannot be excluded.

### Notch signaling strength differs between IFM and DFM progenitors

The distinct effects of Notch depletion on IFM and DFM development prompted us to investigate whether Notch signaling differs between their respective progenitor populations, which can be distinguished by their distinct spatial positions (Figure 3A and C). To assess Notch activity in these populations, we used the Notch-responsive transcriptional reporter *Notch[NRE]-GFP*, which contains a Notch-responsive enhancer (NRE) that drives GFP expression in response to pathway activation [26]. Quantification of *Notch[NRE]-GFP* fluorescence intensity revealed significantly higher reporter levels in IFM AMPs compared with DFM AMPs (Figure 3 A-B). To further assess differences in Notch signaling level, we analysed a second Notch-responsive reporter, *m6-GFP*, whose expression is normally restricted to a subset of IFM AMPs (Figure 3 C-C’). Consistent with *m6-GFP* being a bona fide Notch reporter, depletion of Notch abolished its expression (Figure 3 D-E and G). Conversely, expression of the constitutively active Notch receptor (NotchΔECD) markedly expanded the m6-GFP-positive domain throughout the AMP population (Figure 3 D-F and G). Importantly, this manipulation also induced *m6-GFP* expression in DFM AMPs, where the reporter is normally undetectable (Figure 3F-F’). Together, these results support the conclusion that IFM progenitors are exposed to higher levels of Notch signaling than DFM progenitors. They further indicate that Notch target genes, such as *E(spl)m6*, respond to Notch signaling in a dose-dependent manner, resulting in their differential expression between these progenitor populations.

**Figure 3.**
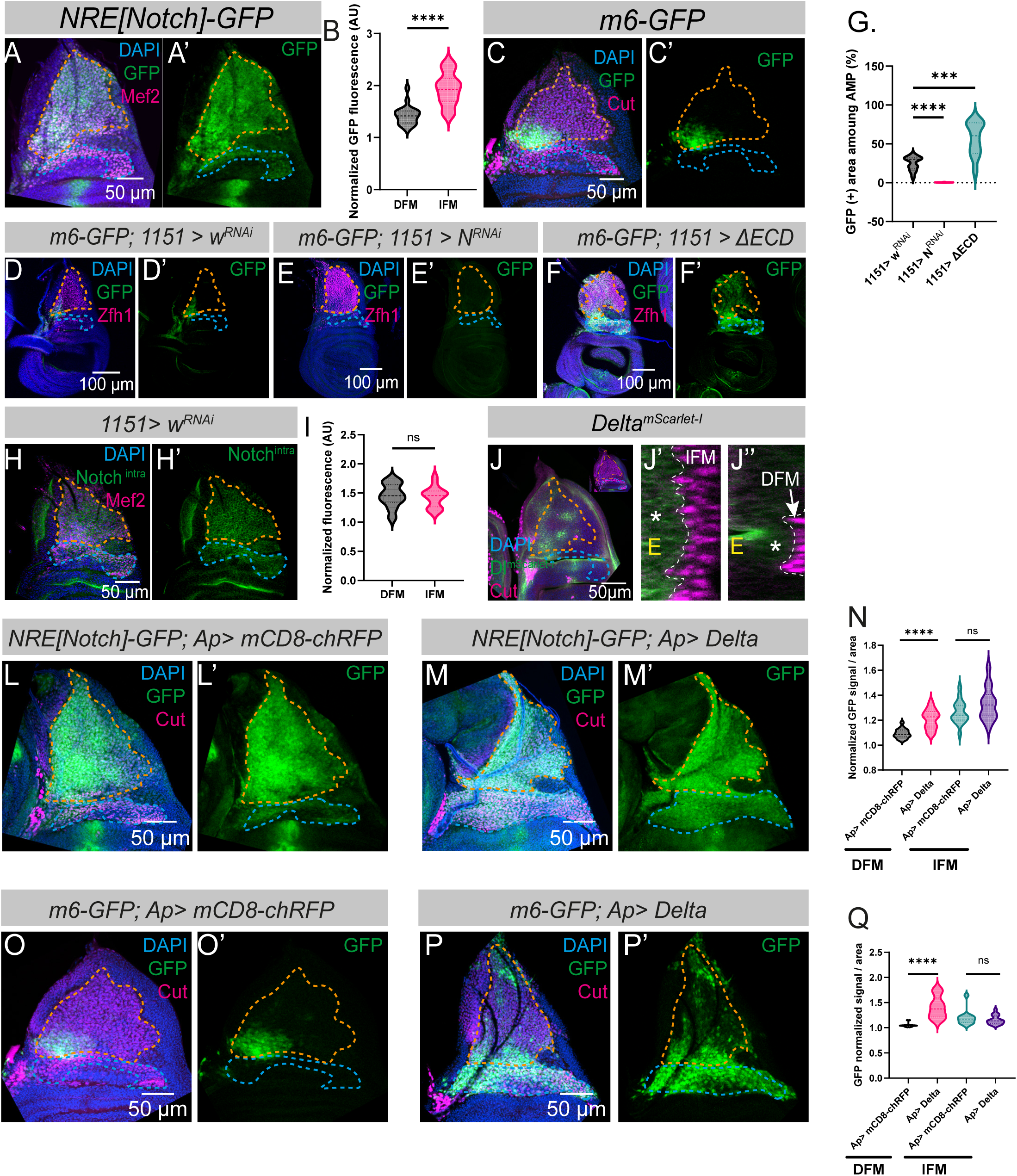
The level of Notch activity differs between direct and indirect flight muscle AMPs and depends on Delta distribution. (A–A′) Expression of the *Notch[NRE]-GFP* reporter in wing discs associated AMPs. GFP (green), Mef2 (magenta) and DAPI (blue). Orange dashed lines delineate the IFM AMPs region, and green dashed lines delineate the DFM AMPs region. **(B)** Quantification of *Notch[NRE]-GFP* fluorescence intensity revealed significantly higher Notch signaling activity in the IFM AMPs than in the DFM AMPs (****, *P* < 0.0001, paired *t*-test; *n* = 22 wing discs from three independent biological replicates). **(C–C′)** Expression of the *E(spl)m6-GFP* reporter (green) in wing disc-associated AMPs. Cut (magenta), GFP (green) and DAPI (blue). **(D–F’)** *E(spl)m6-GFP* expression in control (*1151 > w^RNAi^*) (D, D′), Notch-depleted (*1151 > N^RNAi^*) (E, E′) and Notch-overactivated (*1151 > ΔECD*) (F, F′) conditions. Wing discs are stained for Zfh1 (magenta), GFP (Green) and DAPI (blue). **(G)** Quantification of the *E(spl)m6-GFP*-positive region revealed a significant reduction following Notch depletion and a significant expansion following Notch overactivation (****, *P* < 0.0001; ***, *P* = 0.0003; Student’s *t*-test; control, *n* = 13 wing discs; N^RNAi^, *n* = 13 wing discs; ΔECD, *n* = 12 wing discs; three independent biological replicates). **(H–H’)** Notch intracellular domain (green) expression in wing disc-associated AMPs. Mef2 (magenta) and DAPI (blue). (I) Quantification of Notch-intra fluorescence intensity revealed no significant difference between the IFM and DFM progenitor regions (ns, P = 0.2989, paired t-test; n = 31 wing discs from three independent biological replicates). **(J-J’’)** Delta^mScarlet-I^ expression (green) in the notal region of the wing disc. AMPs are labelled with Cut (magenta) and nuclei with DAPI (blue). **(J’, J’’)** Orthogonal views in the IFMs and DFMs regions. Asterisks indicate enrichment of Delta^mScarlet-I^ adjacent to the IFM progenitor region in (J′), but not adjacent to the DFM progenitor region in (J’’). **(L–M’)** Expression of the *Notch[NRE]-GFP* reporter (green) in wing disc-associated AMPs under control (*Ap > mCD8:RFP*) (L, L’) and epithelial Delta overexpression (*Ap > Delta*) (M, M’) conditions. Wing discs are stained for GFP (green), Cut (magenta) and DAPI (blue). **(N)** *Notch[NRE]-GFP* expression is increased following epithelial Delta overexpression (****, P <0.0001; ns, ***P*** = 0.1174; Student’s *t*-test and Paired *t*-test; control, *n* = 17 wing discs; Delta, *n* = 14 wing discs; three independent biological replicates). **(O–P′)** Expression of the *E(spl)m6-GFP* reporter (green) in wing disc-associated AMPs under control (O, O′) and epithelial Delta overexpression (P, P′) conditions. Wing discs are stained for Cut (magenta) and DAPI (blue). **(Q)** *E(spl)m6-GFP* expression increases following epithelial Delta overexpression (****, *P* < 0.0001; ***, *P* < 0.001; ns; P = 0,4648; Student’s ***t***-test and Mann–Whitney tests; control, *n* = 13 wing discs; *ap > Delta*, *n* = 18 wing discs; two independent biological replicates).

We next sought to determine how this signaling difference is established. To do so, we first examined the distribution of the Notch receptor itself and found that it was homogeneously expressed in IFM and DFM AMPs (Figure 3H-H’ and I), indicating that differential receptor abundance is unlikely to account for the differences in Notch activity. Notch activity in AMPs is induced by the ligand Delta expressed in the adjacent wing disc epithelium [18]. In particular, epithelial cells expressing high levels of Delta are positioned adjacent to the m6-GFP-positive IFM AMPs. Using an endogenously tagged Delta line (*Delta^mScarlet-I^*), we confirmed Delta expression in epithelial cells adjacent to IFM AMPs (Figure 3J–J′). We further found that the Delta signal was stronger in these epithelial cells than in those adjacent to DFM AMPs (Figure 3J’–J’’). If differences in Delta distribution underlie the differential activation of Notch, uniform Delta expression throughout the wing disc epithelium should alter the differential Notch signaling between the two progenitor populations. Consistent with this prediction, epithelial overexpression of Delta strongly increased *Notch[NRE]-GFP* and induced *m6-GFP* expression in DFM AMPs compared with controls (Figure 3L–Q and Figure S1J–K′). Together, these results show that IFM progenitors are exposed to stronger Notch signaling than DFM progenitors and indicate that differential epithelial Delta levels contribute to establishing this difference.

### Differential Notch signaling levels contributes to the distinct Cut and Zfh1 expression in IFM and DFM progenitors

The distinct levels of Notch signaling in DFM and IFM AMPs raise the possibility that quantitative differences in pathway activity influence the expression levels of target genes in these two progenitor populations. One candidate target is the transcription factor (TF) Cut, which is expressed at high levels in DFM progenitors and at lower levels in IFM progenitors (Figure 4A [13]). This expression pattern is opposite to that of Notch signaling, which is higher in IFM than in DFM progenitors. To determine whether differential Notch signaling contributes to the distinct levels of Cut expression, we first depleted Notch in AMPs (Figure 4A-B’). While Notch depletion did not significantly alter Cut fluorescence intensity in DFM progenitors, it induced a modest increase in IFM progenitors, although this increase did not reach statistical significance (Figure 4C-D). We therefore asked whether increasing Notch signaling was sufficient to repress Cut expression. To this end, we expressed a constitutively active form of the Notch receptor (*Notch*Δ*ECD*) throughout the AMP population using *1151-Gal4*. NotchΔECD significantly reduces Cut expression in both DFM and IFM progenitors (Figure 4 E-F and G-H). Given that Delta is normally enriched in the wing disc epithelium adjacent to IFM progenitors, we next asked whether increasing Delta levels uniformly throughout the epithelium would similarly equalize Cut expression between the two progenitor populations (Figure S1 J-K’). Consistent with the effects of *Notch*Δ*ECD*, epithelial Delta overexpression abolished the difference in Cut expression between DFM and IFM progenitors (Figure 4 I-J’ and K-L). Together, these results indicate that Notch negatively regulates Cut expression in AMPs and suggest that the lower level of Notch signaling in DFM progenitors contributes to maintaining their higher Cut expression.

**Figure 4.**
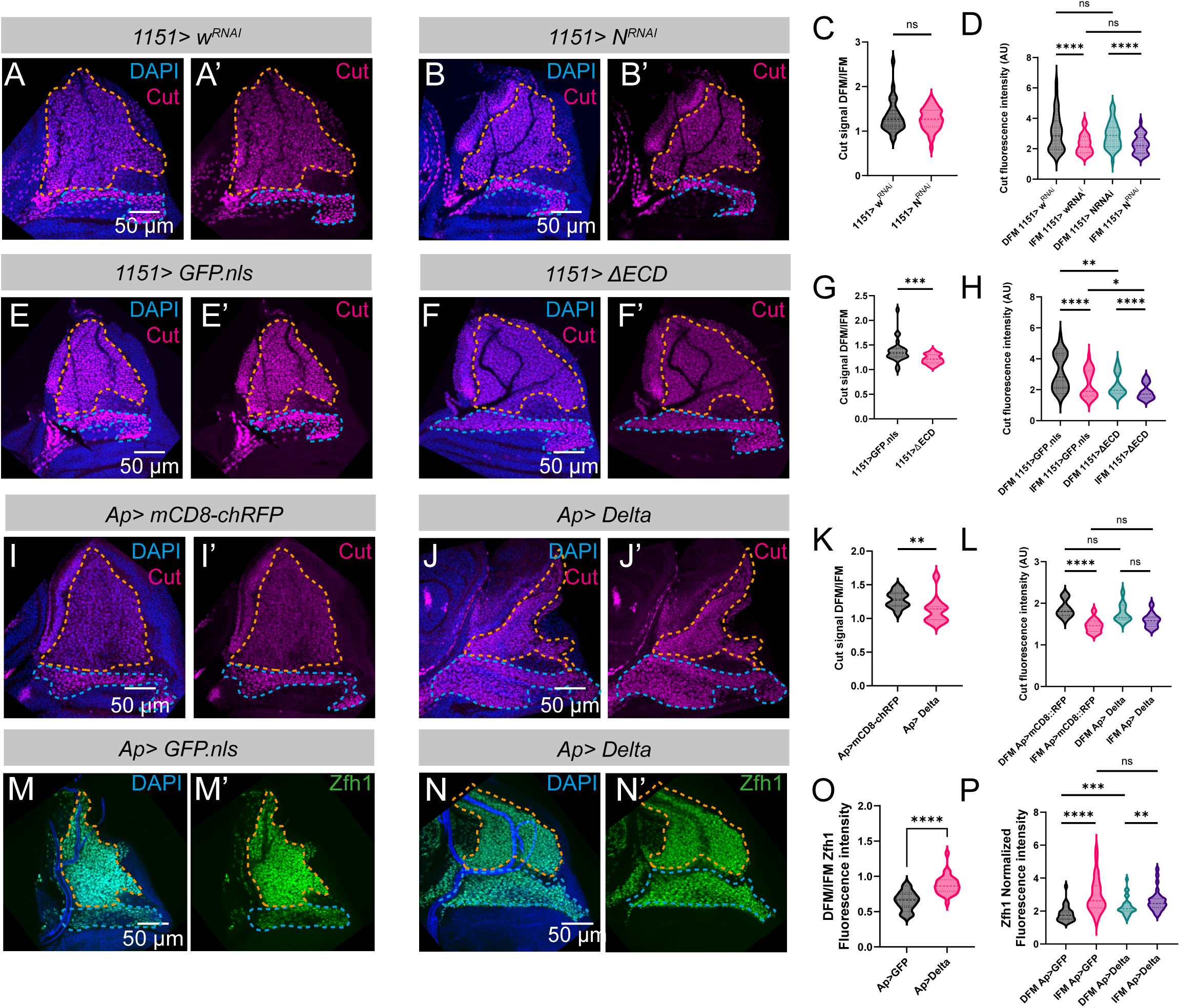
Notch signaling contributes to differential Cut and Zfh1 expression between IFM and DFM progenitors. (A–B′) Wing discs stained for Cut (magenta) and DAPI (blue) under control (*1151 > w^RNAi^*) (A, A′) and Notch-depleted (*1151 > N^RNAi^*) (B, B′) conditions. **(C)** Quantification of the Cut fluorescence intensity ratio between the IFM and DFM AMPs regions revealed no significant difference following Notch depletion (ns, *P* = 0.9552; Mann–Whitney test; control, *n* = 33 wing discs; N^RNAi^, *n* = 39 wing discs; three independent biological replicates). **(D)** Quantification of Cut fluorescence intensity revealed no significant change following Notch depletion in either the IFM or DFM AMPs regions (**, *P* = 0.0068; ***, *P* = 0.0008; Mann–Whitney tests; control, *n* = 33 wing discs; N^RNAi^, *n* = 39 wing discs; three independent biological replicates). **(E–F′)** Wing disc stained for Cut (magenta) and DAPI (blue) under control (*1151 > w^RNAi^*) (E, E′) and Notch-overactivated (*1151 > ΔECD*) (F, F′) conditions. **(G)** Quantification of the Cut fluorescence intensity ratio between the IFM and DFM progenitor regions revealed a significant reduction following Notch overactivation (**, *P* = 0.0003; Mann– Whitney test; control, *n* = 27 wing discs; ΔECD, *n* = 25 wing discs; four independent biological replicates). **(H)** Quantification of Cut fluorescence intensity revealed a significant reduction in both the IFM and DFM AMPs regions following Notch overactivation (****, *P* < 0.0001; ***, *P* < 0.001; **, *P* < 0.01; *, *P* < 0.05; Mann–Whitney and Wilcoxon tests; control, *n* = 27 wing discs; ΔECD, *n* = 25 wing discs; four independent biological replicates). **(I–J′)** Wing disc stained for Cut (magenta) and DAPI (blue) under control (*Ap > mCD8:chRFP*) (I, I′) and epithelial Delta overexpression (*Ap > Delta*) (J, J′) conditions. **(K)** Cut fluorescence intensity ratio between the IFM and DFM progenitor regions is reduced following epithelial Delta overexpression (***, *P* = 0.0086; Mann–Whitney test; control, *n* = 12 wing discs; Delta *n* = 11 wing discs; two independent biological replicates). **(L)** Quantification of Cut fluorescence in the IFM and DFM AMPs in control and following epithelial Delta overexpression (***, *P* = 0.0005; Student’s *t*-test, Paired t test, Mann–Whitney and Wilcoxon tests; control, *n* = 12 wing discs; Delta *n* = 11 wing discs; two independent biological replicates). **(M–N′)** Wing discs stained for Zfh1 (green) and DAPI (blue) under control (*Ap>GFP.nls*) (M-M’) and epithelial Delta overexpression (*Ap > Delta*) (N-N’). **(O)** Zfh1 fluorescence intensity ratio between the DFM and IFM progenitors regions is increased following epithelial Delta overexpression (****, <0,0001; Student t test; control, *n* = 28 wing discs; Delta *n* = 29 wing discs; two independent biological replicates). **(P)** Zfh1 fluorescence is significantly increased in DFM progenitors following epithelial Delta overexpression (****, *P* < 0.0001; ***, P = 0.0002; **, = 0.0091; ns, P = 0.4049; Mann–Whitney or Wilcoxon tests; control, *n* = 28 wing discs; Delta *n* = 29 wing discs; two independent biological replicates).

We next examined the TF Zfh1, a known direct Notch target in AMPs [20, 27]. In contrast to Cut, Zfh1 is expressed at higher levels in IFM progenitors and at lower levels in DFM progenitors (Figure 4M-P). This pattern mirrors the distribution of *Notch[NRE]-GFP* reporter expression, suggesting that differential Notch signaling driven by Delta may contribute to establishing distinct Zfh1 levels between the two progenitor populations. To test this hypothesis, we overexpressed Delta in the wing disc epithelium and found that Zfh1 expression significantly increased in DFM progenitors (Figure 4N–P). Together, these results indicate that quantitative differences in Notch signaling contribute to distinct gene expression levels in IFM and DFM progenitors. This raises the question of how these differences in signaling are decoded into distinct transcriptional responses.

### IFM and DFM progenitors have distinct transcriptional bursting dynamics in response to Notch signaling

Changes in transcriptional burst dynamics can generate different levels of gene expression [28]. We therefore asked whether the distinct levels of Notch signaling in IFM and DFM AMPs are translated into differences in transcriptional bursting dynamics that, in turn, contribute to their distinct gene expression profiles. In order to monitor transcription in living cells, we utilized the MS2-MCP system [29, 30]. We generated a Notch transcriptional sensor, where transcription of a yellow transcription unit is controlled by the Notch-responsive enhancer (NRE) (Figure 5A). In order to maximize signal detection, we inserted 24 MS2 in the 5’UTR region, as designed in embryonic reporters [31] (Figure 5A). The specificity of the reporter expression was first validated in fixed tissues by single-molecule fluorescence in situ hybridization (smFISH) using a probe against the *yellow* sequence. *Notch-MS2* transcripts were specifically detected in the AMP population (Figure 5B–C′) and precisely overlapped with the *Notch[NRE]-GFP* reporter (Figure S2A). We next assessed reporter responsiveness by expressing NotchΔECD in AMPs and using smFISH to detect nascent transcripts at transcription sites (TS) [32, 33]. Our quantification revealed a significant increase in the proportion of transcriptionally active nuclei upon overexpression of the Notch pathway (Figure S2 A-C). Together, these experiments validate *Notch-MS2* as a sensitive reporter for monitoring dynamic Notch-dependent transcription in living AMPs. The *Notch-MS2* reporter was then combined with *MCP-GFP* and *His2Av-mRFP* to simultaneously visualize TS and track individual nuclei over time. Live imaging of isolated living wing discs revealed bright MCP-GFP puncta within AMP nuclei (Figure D-D’ and Movie S1). These puncta precisely colocalized with nascent *yellow* transcription foci detected by smFISH (Figure S2 E-F’’), confirming that they represent bona fide transcription sites of the *Notch-MS2* reporter.

**Figure 5.**
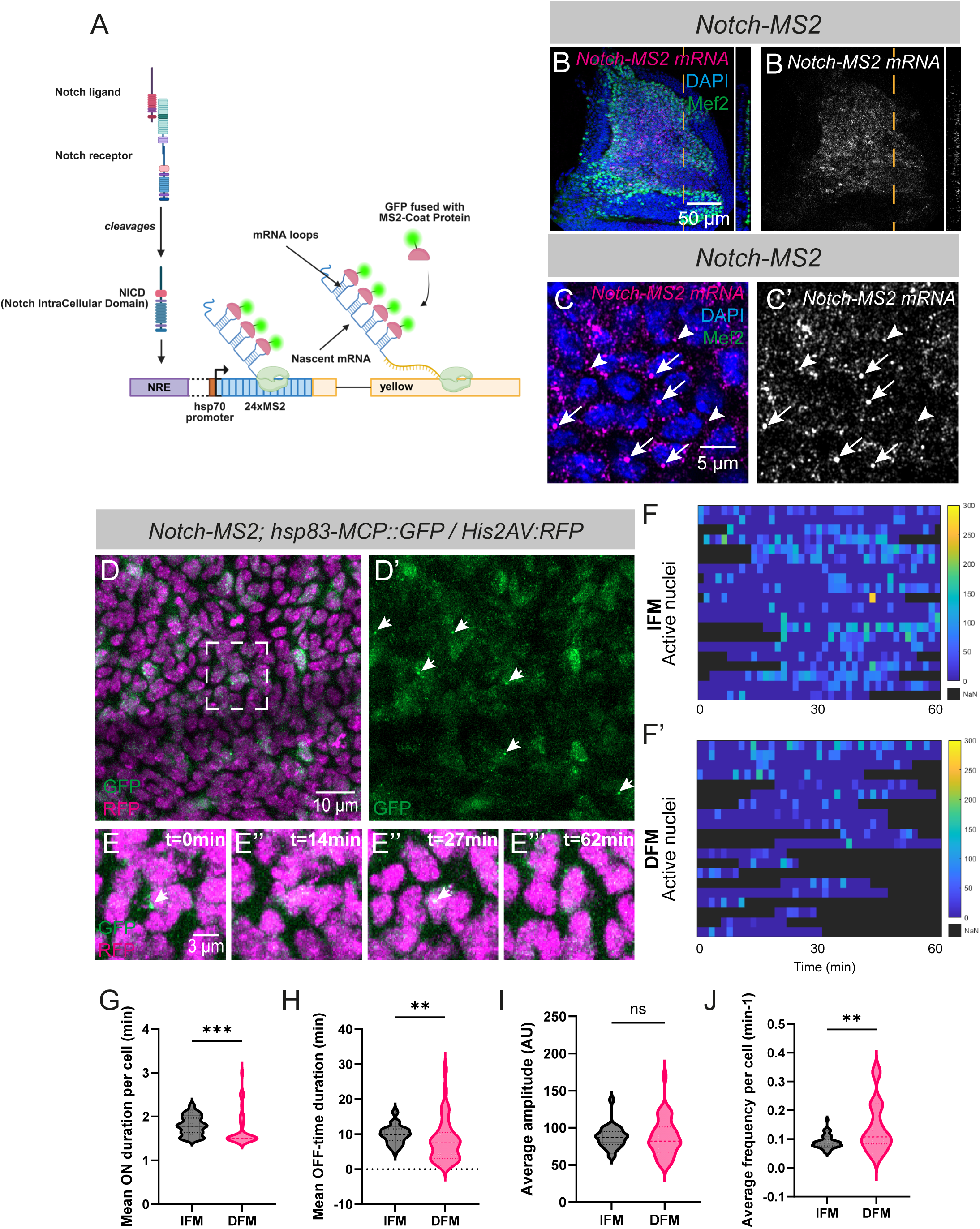
Notch-dependent transcriptional bursting dynamics differ between IFM and DFM progenitors. **(A)** Schematic representation of the *Notch-MS2* reporter. The Notch Responsive Element (NRE) is inserted upstream of the MS2 repeats, followed by a truncated *yellow* gene. During transcription, the free MCP-GFP binds to the MS2 stem loops, allowing visualization of active transcription sites. **(B–C’)** Detection of *Notch-MS2* transcripts (grey) by smFISH with a *yellow* probe in wing disc-associated AMPs (Mef2, green). **(C–C’)** Higher-magnification image shows *Notch-MS2* active transcription sites (arrowheads) and individual mRNA molecules (arrows). **(D-D’)** Single time point from a time-Notch-MS2 lapse sequence (see Movie S1) showing *Notch-MS2* transcription (green) in wing disc-associated AMPs. Nuclei are labelled with His2AV::RFP (magenta). Arrowheads indicate nuclei undergoing active transcription of the *Notch-MS2* reporter. **(E–E’’’)** Time course (min) of the nuclei outlined in (D), showing detection of the active *Notch-MS2* transcription site over time. **(F–F′)** Heatmaps showing *Notch-MS2* transcriptional activity over a 60-min period in transcriptionally active IFM (F) and DFM (F′) AMP nuclei. Fluorescence intensity is represented by the colour scale, with consecutive time points separated by 1 min 30 s. **(G–J)** Quantification of transcriptional bursting parameters revealed significantly longer burst ON durations (G) and OFF durations (H) in IFM AMPs compared with DFM AMPs. Burst amplitude did not differ significantly between the two populations (I), whereas burst frequency was significantly higher in DFM AMPs (J). (**, *P* = 0.0017; Mann–Whitney test; DFM n = 39 nuclei; IFM n = 53 nuclei; N = 3 movies of each population from three independent biological replicates).

As an additional specificity control, expression of MCP-GFP alone produced diffuse fluorescence throughout the AMP population with no detectable nuclear puncta (Figure S2G–G′′′). In contrast, the presence of the *Notch-MS2* reporter resulted in the appearance of discrete nuclear foci (Figure S2H–H′′′), consistent with MCP-GFP binding to nascent MS2-containing transcripts. Analysis of the movies showed that TS detection was highly heterogeneous and asynchronous between nuclei (Movie S1). Individual TS dynamically appear and disappear over time (Figure 5 E-E’’’), pointing to their discontinuous activity. These observations are consistent with the smFISH analysis, which revealed variability in TS detection among AMPs (Figure 5C). This initial analysis revealed that Notch-dependent transcription in the AMPs occurs in bursts.

Individual nuclei were then tracked over time, and the fluorescence intensity of each TS was extracted to generate temporal transcriptional profiles for IFM and DFM AMPs (Figure 5F–F′). These profiles were used to quantify the four principal parameters of transcriptional bursting: burst ON duration (the time during which transcription is active), burst OFF duration (the interval between successive bursts), burst amplitude (the maximal transcriptional output during a burst), and burst frequency (the number of bursts per unit time). Our analysis revealed that IFM progenitors displayed significantly longer burst ON durations than DFM progenitors (Figure 5G). Burst OFF durations were also significantly different between the two populations, with longer median OFF periods in IFM progenitors (Figure 5H). In contrast, burst amplitude did not differ significantly between IFM and DFM progenitors (Figure 5I), whereas burst frequency was significantly higher in DFM progenitors (Figure 5J).

We next performed smFISH to determine whether these differences in transcriptional dynamics were reflected in the proportion of transcriptionally active nuclei. We found that a significantly larger fraction of IFM progenitors contained detectable *Notch-MS2* TS than DFM progenitors (Figure 6A–C). Similarly, the proportion of nuclei actively transcribing *zfh1* was higher in IFM than in DFM progenitors (Figure 6D–F). This difference is reminiscent of the differential levels of Zfh1 protein observed between IFM and DFM progenitors (Figure 4M–P).

**Figure 6.**
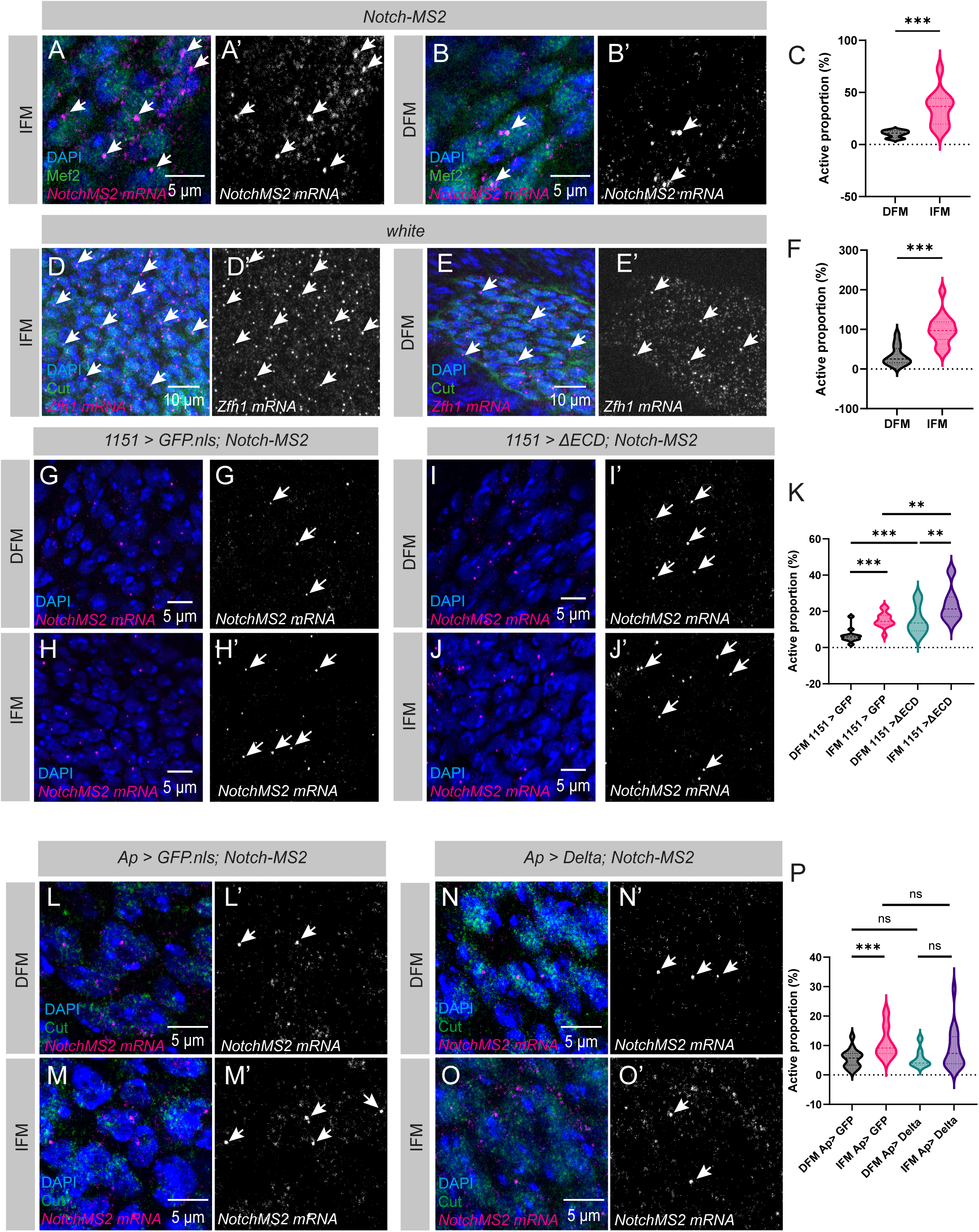
Differential Notch-dependent transcription in IFM and DFM progenitors is regulated by Delta. (A–B′) Detection of active Notch-MS2 transcription sites by smFISH in the IFM (A, A′) and DFM (B, B′) AMPs. Mef2 is shown in green and Nuclei (DAPI) in blue. **(C)** Quantification of the proportion of transcriptionally active nuclei revealed a higher proportion of active cells in the IFM AMPs than in the DFM AMPs (***, *P* = 0.0003; Student’s *t*-test; DFM, n = 796 nuclei; IFM n = 1449; *N* = 12 wing discs from three independent biological replicates). **(D–E′)** Detection of active zfh1-long transcription sites by smFISH in the IFM (D, D′) and DFM (E, E′) AMPs. (**F**) Quantification of the proportion of transcriptionally active nuclei revealed a higher proportion of active cells in the IFM AMPs than in the DFM AMPs (***, *P* = 0.0005; Wilcoxon test; DFM, n = 2144 nuclei; IFM n = 4761; *N* = 16 wing discs from three independent biological replicates). (**G-J’**) Detection of active Notch-MS2 transcription sites by smFISH in the DFM (G-G’, I-I’) and IFM (H-H’, J-J’) AMPs under control (*1151 > GFP.nls*) (G-H’) and overactivated Notch (*1151 > ΔECD*) (I-J’) conditions. **(K)** The proportion of the nuclei actively transcribing Notch-MS2 is increased in both the IFM and DFM progenitor regions following Notch overactivation (****, *P* < 0.0001; ***, *P* < 0.001; **, *P* < 0.01; Student’s *t*-test, Mann– Whitney or Wilcoxon tests; control, *n* = 12 wing discs; ΔECD, *n* = 13 wing discs; four independent biological replicates). **(L-O’)** Detection of Notch-MS2 transcription by smFISH in DFM (L-L’, N-N’) and IFM (M-M’, O-O’) progenitors under control (Ap> GFP.nls) (L-M’) and Delta overexpression (*Ap> Delta*) (N-O’) conditions. Nuclei are stained with Cut (Green) and DAPI (blue). Notch-MS2 transcripts are shown in magenta and grey. **(P)** Epithelial Delta overexpression equalized the proportion of nuclei actively transcribing the Notch-MS2 reporter between DFM and IFM AMPs. (*, *P* = 0.0159; Mann–Whitney test; control, *n* = 15 wing discs; Delta, *n* = 15 wing discs; four independent biological replicates).

Together, these findings show that IFM and DFM progenitors have distinct dynamics of Notch-dependent transcription. IFM progenitors display longer transcriptional bursts and a higher proportion of transcriptionally active nuclei compared with DFM progenitors. Such differences in transcriptional dynamics may contribute to the higher expression of Notch-activated genes, such as *zfh1*, in IFM progenitors, while lower Notch signaling in DFM progenitors may favor the expression of negatively regulated genes, such as *cut*.

### The spatial distribution of epithelial Delta shapes Notch-dependent transcription in IFM and DFM progenitors

The finding that IFM and DFM progenitors show distinct Notch-dependent transcriptional responses raised the question of whether these differences result from the different levels of Notch signaling received by the two populations. To address this, we first expressed a constitutively active form of Notch in AMPs using the *1151-Gal4* driver and compared the proportion of transcriptionally active nuclei between IFM and DFM progenitors using the *Notch-MS2* reporter. Notch overactivation significantly increased the proportion of transcriptionally active nuclei in both populations, while reducing the difference between them (Figure 6G–K).

We next asked whether the spatial distribution of Delta in the epithelial niche contributes to this differential transcriptional response. We therefore overexpressed Delta throughout the wing disc epithelium using *Ap-Gal4* and quantified the proportion of transcriptionally active nuclei in both populations. While IFM progenitors displayed a higher proportion of active nuclei than DFM progenitors under control conditions, this difference was abolished following epithelial Delta overexpression, resulting in comparable proportions of transcriptionally active nuclei in the two populations (Figure 6L–P). Together, these results indicate that the spatial distribution of Delta in the epithelial niche contributes to establishing the differential Notch-dependent transcriptional responses of IFM and DFM progenitors.

## Discussion

Here, we investigated how quantitative differences in Notch signaling contribute to the specification of distinct muscle progenitor populations. We show that IFM and DFM progenitors differ in their levels of Notch signaling and that, despite the relatively low pathway activity in DFM progenitors, Notch remains required for their proper development. We further demonstrate that the spatial distribution of Delta in the wing disc epithelium contributes to these quantitative differences, which are associated with distinct expression patterns of Notch-regulated genes. Using live transcriptional imaging, we reveal that Notch-dependent transcription in AMPs occurs in bursts, with IFM progenitors displaying longer but less frequent bursts than DFM progenitors. Together, our findings suggest that spatial differences in ligand availability contribute to quantitative differences in Notch signaling, which may be translated into distinct transcriptional responses through modulation of transcriptional bursting dynamics.

It is now widely accepted that gene transcription occurs in stochastic bursts in both prokaryotic and eukaryotic cells [8, 34–36]. However, only a limited number of studies have directly examined how developmental signaling pathways regulate transcriptional bursting *in vivo*. Two pioneering studies employed the MS2 system to investigate the transcriptional dynamics of the Notch response [10, 11]. Lee et al. showed that Notch-dependent transcription in *C. elegans* germline stem cells occurs in stochastic bursts and, importantly, that increased Notch signaling preferentially extends transcriptional burst duration. Independently, Falo-Sanjuan et al. used a similar approach to monitor Notch-responsive enhancers in the early *Drosophila* embryo, and revealed that Notch target genes are likewise transcribed in stochastic bursts and that increased Notch levels alters the amplitude and duration of the transcription. Our findings extend these studies in several important ways. First, we show that stochastic transcriptional bursting also underlies the Notch response during adult muscle progenitor specification, a developmental process unfolding over a much longer timescale than embryogenesis. More importantly, we described a physiological developmental context in which neighbouring muscle progenitors are exposed to distinct doses of endogenous Notch signaling. We show that these quantitative signaling differences are decoded through modulation of transcriptional burst duration. Finally, we identify the spatially restricted distribution of Delta as the source of these quantitative differences in Notch signaling.

How modulation of transcriptional burst generates distinct developmental outcomes remains an important and largely unresolved question. Most studies investigating transcriptional bursting in *Drosophila* have been performed during early embryogenesis, where cells are still largely undifferentiated and developmental outcomes cannot be readily linked to transcriptional dynamics. However, recent work in the developing *Drosophila* eye provided evidence that transcriptional burst dynamics can directly influence cell fate by controlling the proportion of distinct cell types produced [37]. In this study, we found that Notch-dependent transcription is characterized by longer burst durations in IFM progenitors than in DFM progenitors. One possible consequence of longer bursts is an increase in the number of transcripts produced during each promoter activation event, thereby increasing overall target gene expression without altering burst frequency. Such quantitative differences could enable Notch target genes with distinct activation thresholds to respond differently to signaling, ultimately generating distinct transcriptional programs and developmental outcomes. This mechanism may be particularly relevant for genes whose expression is graded rather than binary. For example, the TF Vestigial and Cut are expressed in both IFM and DFM progenitors but at different levels, with Vestigial enriched in IFM progenitors and Cut enriched in DFM progenitors [13]. It is therefore tempting to speculate that differences in Notch-dependent transcriptional burst duration contribute to establishing these expression thresholds. In IFM progenitors, prolonged Notch-dependent transcription may cooperate with Wingless signaling to promote high Vestigial expression while limiting Cut expression. Conversely, the shorter transcriptional bursts observed in DFM progenitors may be insufficient to activate the same transcriptional program, thereby allowing Cut to remain highly expressed.

Notch-responsive enhancers may act as molecular decoders of quantitative differences in signaling. Their cis-regulatory architecture, including the number, affinity and organization of Su(H)-binding sites, the integration of additional transcriptional inputs such as Wingless signaling, and the local chromatin environment, may determine how individual Notch target genes translate signaling levels into distinct transcriptional bursting dynamics. Such regulation could allow neighbouring progenitors exposed to different levels of Notch signaling to activate overlapping sets of target genes while producing distinct levels of gene expression. Understanding how quantitative Notch signaling is interpreted by cis-regulatory elements and translated into transcriptional bursting dynamics will therefore be essential to uncovering how a single signaling pathway contributes to the specification of distinct cell identities.

## Materials and Methods

### Drosophila melanogaster strains and genetics

All *Drosophila melanogaster* stocks were grown on standard medium at 25°C. The following stains were used: *w^1118^* as wild type (wt), *UAS-w^RNAi^* (BL#28980), *UAS-N^RNAi^* (BL#7078), *UAS-Notch*Δ*ECD* [38–40], *UAS-Delta^RNAi^*(VDRC#109461), *UAS-Delta* [41], *1151-Gal4* [22], *Ap-Gal4* (BL#3041), *Delta^mScarlet-^*^I^ [18], *UAS-mCD8-chRFP* (BL#27392), *m6-GFP* [42], *NRE[Notch]-GFP* [26], *UAS-GFP.nls* (BL#4776), *Hsp83-MCP-GFP* (BL#7280), *UAS-MCP-GFP* (VDRC# 311057), *His2Av-mRFP* (BL#23650).

### Generation of the *Notch-MS2* fly line

The Notch Responsive Element (NRE), corresponding to the enhancer used in the *Notch[NRE]-GFP* reporter line [26], was cloned upstream of the 24×MS2-*yellow* reporter cassette [31] using the NotI and BamHI restriction sites. The resulting Notch-MS2 construct was integrated into the attP51D landing site on chromosome II following embryo injection into *vas-int; attP51D* flies.

### Immunostainings and in situ hybridization

Wing disc and adult muscle preparations, immunofluorescence, and single-molecule fluorescence in situ hybridization (smFISH) were performed as previously described [33]. The following primary antibodies were used for immunofluorescence staining: chicken anti-GFP (1:300; Abcam, ab13970), rabbit anti-Mef2 (1:500; DSHB, Mef2-p), rabbit anti-Zfh1 (1:5,000; a gift from Ruth Lehmann, New York University, USA), mouse anti-MHC (1:200; DSHB, 3E8-3D3-s), mouse anti-Cut (1:10; DSHB, 2B10), mouse anti-Delta (1:50; DSHB, C594.9B), and mouse anti-Notch intracellular domain (NICD) (1:100; DSHB). Nuclei were counterstained with DAPI.

### Flight assay

Flight ability was assessed using a 1-m-long cylindrical flight column internally lined with paper coated with a thin layer of glue, similarly as in (Babcock & Ganetzky 2014). The adhesive sheet was divided evenly into three longitudinal sections corresponding to the Top, Middle, Bottom and Ground regions of the tube. For each experimental condition, an equal number of flies (approximately 30) was released into the tube. After all flies had landed, the paper was removed, flattened, and imaged for analysis.

Image quantification was performed semi-automatically using a custom R script, Fly Spy (available at https://github.com/Olivier-Josse/Fly-spy). Images were first converted to grayscale and digitally divided into four predefined regions. High intensity spots corresponding to individual flies were identified by applying a manually adjusted threshold to ensure accurate segmentation. The number of flies detected in each region was quantified and expressed as a percentage of the total recovered flies. The distribution of flies across the four regions—reflecting high, intermediate, low or absent flight ability—was compared between experimental conditions using a chi square (χ²) test. Results are presented as stacked bar histograms.

### Microscopy and data analysis

#### Fixed samples

Fluorescent fixed samples were imaged at the BIOSIT Imaging Facility (University of Rennes) using either a Zeiss LSM980 Airyscan 2, a Leica SP8, or a Nikon N-SPARC confocal microscope equipped with 20×, 40×, or 63× objectives. Images were processed using Fiji/ImageJ, and final figures were prepared in Adobe Illustrator. For fluorescence measurements, all genotypes were processed, imaged, and analyzed in parallel. Within each experiment, identical acquisition settings (laser power and detector gain) were used for all samples. Sum-intensity projections were generated from each confocal z-stack before analysis. Regions of interest (ROIs) were manually delineated according to the expression pattern of the marker being quantified, and the mean fluorescence intensity was measured for each ROI. Fluorescence values were normalized to background intensity measured in an adjacent region of the same sample.

#### Quantification of myonuclear density and sarcomere length

For myonuclear density analysis, nuclei were counted manually and normalized to the area of the analyzed muscle fiber. Sarcomere length was measured using the ImageJ plugin described in [43]. A line was drawn along individual myofibrils, and the length of each sarcomere was determined. Three myofibrils were analyzed per muscle, and measurements were pooled to calculate the mean sarcomere length for each muscle.

#### Live imaging

Larvae were dissected in Schneider’s Drosophila medium, as described in [44]. Wing discs were mounted in Schneider’s medium supplemented with 0.6% penicillin–streptomycin (Invitrogen 15140-122), 15% fetal bovine serum (Biosera FB-1200/500), and insulin (Sigma I5500). Samples were mounted between an 18 × 18 mm coverslip (VWR 631-1567) and a 35 mm Lumox dish. To prevent evaporation during acquisition, preparations were covered with Halocarbon oil 27 (Sigma H8773). Live imaging was performed on an LSM980 Airyscan2 microscope (BIOSIT, University of Rennes) with the chamber maintained at 25°C. Time-lapse sequences were acquired using a 63× objective, at a resolution of 480 × 480 pixels, with images collected every 90 seconds.

#### Single-molecule FISH analysis

Transcriptional spots were detected and counted by an ImageJ script using Trackmate plugin in Fiji/ImageJ, in 3D, as previously described [33]. Due to variations in fluorescence intensity among samples, “radius” and “quality” parameters were manually selected specifically for each sample. Different regions of the same sample were analyzed with the same parameters (as example IFM and DFM regions). Nuclei were counted manually in the total volume. To obtain the active proportion, total number of spots was divided by the total number of nuclei.

#### MS2 data processing and quantitative analysis

Nuclei were detected and segmented in 3 dimensions (3D) using Cellpose [45] and tracked over time using the TrackMate plugin in Fiji/ImageJ [46], with a maximum linking distance of 2 µm between consecutive frames. An adapted MATLAB script (MATLAB 2025b, MathWorks) was used to detect and track NotchMS2 transcription foci [11]. Transcription foci were detected in 3D. Heatmaps were generated for each movie to visualize the temporal transcriptional activity of nuclei exhibiting at least one detectable transcriptional burst during the recording. For each tracked nucleus, we automatically quantified the duration of transcriptionally active (ON) and inactive (OFF) periods, burst amplitude (maximum normalized transcription site intensity), and burst frequency (number of detected burst cycles per unit of recording time). To avoid including incomplete events at the beginning or end of recordings, ON periods were retained only when flanked by two OFF periods. Similarly, OFF periods were retained only when flanked by two ON periods. A maximum gap of two consecutive frames without nuclear detection was tolerated during OFF periods. Nuclei that did not meet the criteria for a given parameter were excluded from the corresponding analysis. To account for variations in fluorescence intensity between nuclei, the intensity of each transcription site was normalized to the mean MCP-GFP fluorescence intensity measured within the corresponding nuclear volume in 3D.

## Statistical analysis

Normality was checked via a Shapiro-Wilk test. When data were normally distributed, a student t-test was applied. Otherwise, a Mann-Whitney test was used. In the case of paired data, for normally distributed data, paired t test was applied, otherwise, Wilcoxon test was applied. In all cases, significance was presented as ****, p<0.0001,***, p<0.001 **, p<0.01, *, p<0.05, and P values are provided in figure legends.

## Acknowledgments

We would like to thank the Bloomington Drosophila Stock Center, the Vienna Drosophila Resource Center (VDRC), and the Developmental Studies Hybridoma Bank for providing *Drosophila* strains and antibodies. We are grateful to Tanguy Lucas, Carmen Santa Cruz Mateos and Alain Vincent for the critical reading of the manuscript. We acknowledge Xavier Pinson and Stephanie Dutertre from the Microscopy Rennes Imaging Center (MRic, BIOSIT, Biogenouest) for technical assistance. We are also grateful to members of the Sarah Bray laboratory for hosting E.L. and providing training in MS2-based transcriptional dynamics analysis during the initial stages of this project. Work in Boukhatmi lab is supported by ATIP-Avenir (CNRS), ANR GenepiMuSC (ANR-23-CE13-0006), and la Ligue contre le Cancer. Work in Lagha lab is supported by the Centre National de la Recherche Scientifique and the Université de Montpellier, and is funded by grants from the European Research Council (SyncDev, LightRNA2Prot) and Agence Nationale de la Recherche HubDyn. E.L was supported by an AFM-Téléthon PhD fellowship (#29992). The funders had no role in study design, data collection and analysis, decision to publish, or preparation of the manuscript.

## Competing interest

The authors declare that they have no competing interests.

---

**Figure S1.**
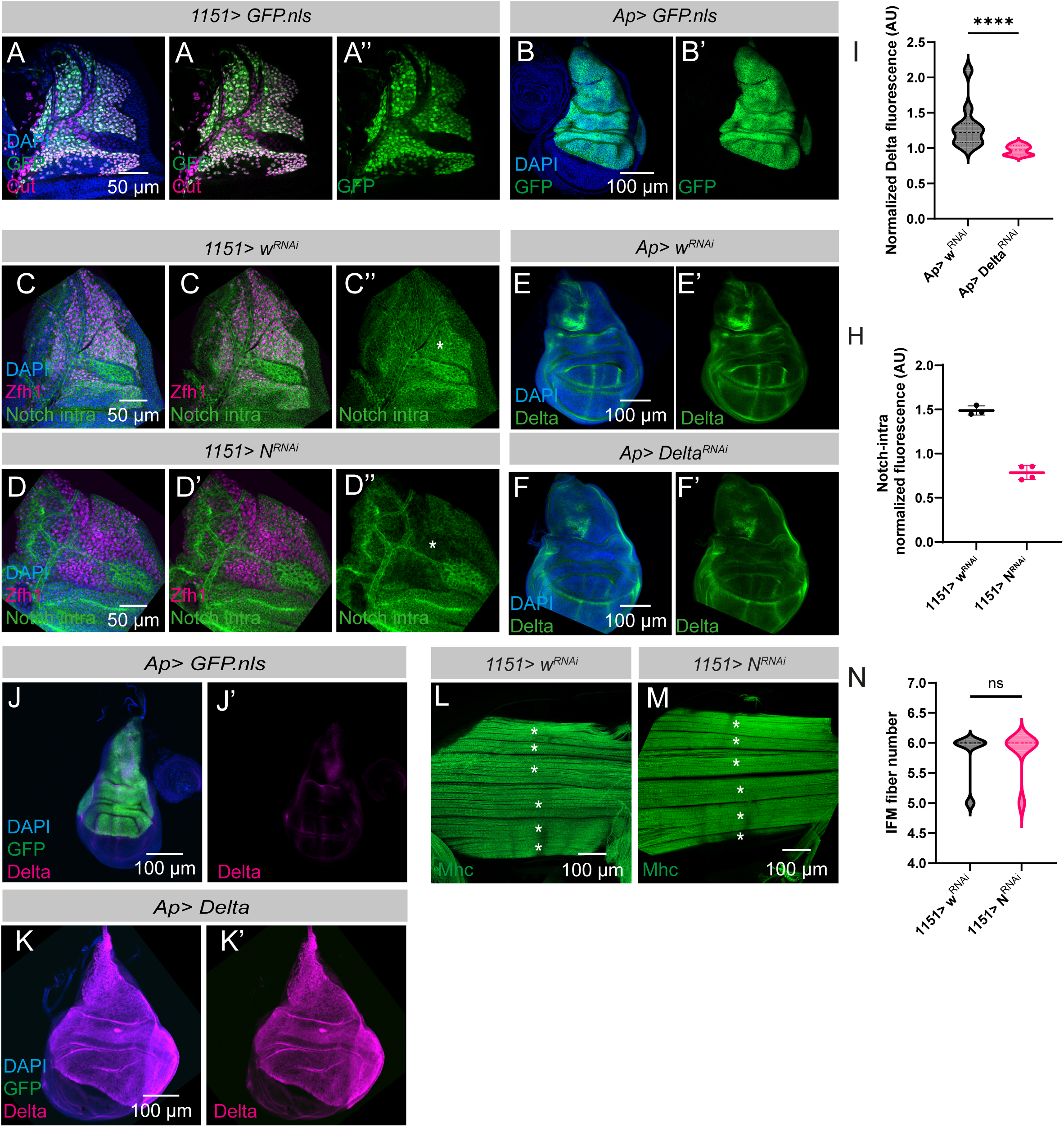
(A–A’′) GFP expression driven by *1151-Gal4* (green) in the wing disc-associated AMPs. AMPs are labelled with Cut (magenta) and nuclei with DAPI (blue). **(B–B′)** GFP expression driven by *ap-Gal4* (green) in wing disc epithelial cells. Nuclei are labelled with DAPI (blue). **(C–D″)** Distribution of the Notch intracellular domain (NICD; green) in control (C–C″) and Notch-depleted (*1151 > N^RNAi^*) (D–D″) AMPs. Zfh1 labels AMPs (magenta) and nuclei are stained with DAPI (blue). **(E)** Quantification of Notch intracellular domain fluorescence intensity revealed a significant reduction following Notch depletion in AMPs. **(F–H’)** Delta expression (green) in wing discs under control (*Ap > w^RNAi^*) (F–F’) and Delta-depleted (*Ap > Delta^RNAi^*) (H– H’) conditions. Delta is detected in the notal region and along the dorsoventral (DV) boundary. **(I)** Quantification of Delta fluorescence intensity revealed significantly lower expression in the epithelial Delta-depleted condition (***, *P* < 0.0001; Mann–Whitney test; control *n* = 11 wing discs; Delta^RNAi^ *n* = 10 wing discs; from two independent biological replicates). **(J-K’)** Wing disc stained for Delta (magenta) and DAPI (blue) under control (*Ap > mCD8:chRFP*) (J-J′) and epithelial Delta overexpression (*Ap > Delta*) (K, K′) conditions. **(L-M)** Indirect flight muscles (IFMs) stained for Mhc (green) in control (*1151 > w^RNAi^*) (L) and Notch-depleted (*1151 > N^RNAi^*) (M) adult flies. (**N**) Quantification of fiber number revealed no significant differences between control and Notch-depleted conditions (ns, *P* = 0.7881; Student t test; control *n* = 10 hemi thoraxes; N^RNAi^ *n* = 13 hemi thoraxes; from three independent biological replicates).

**Figure S2.**
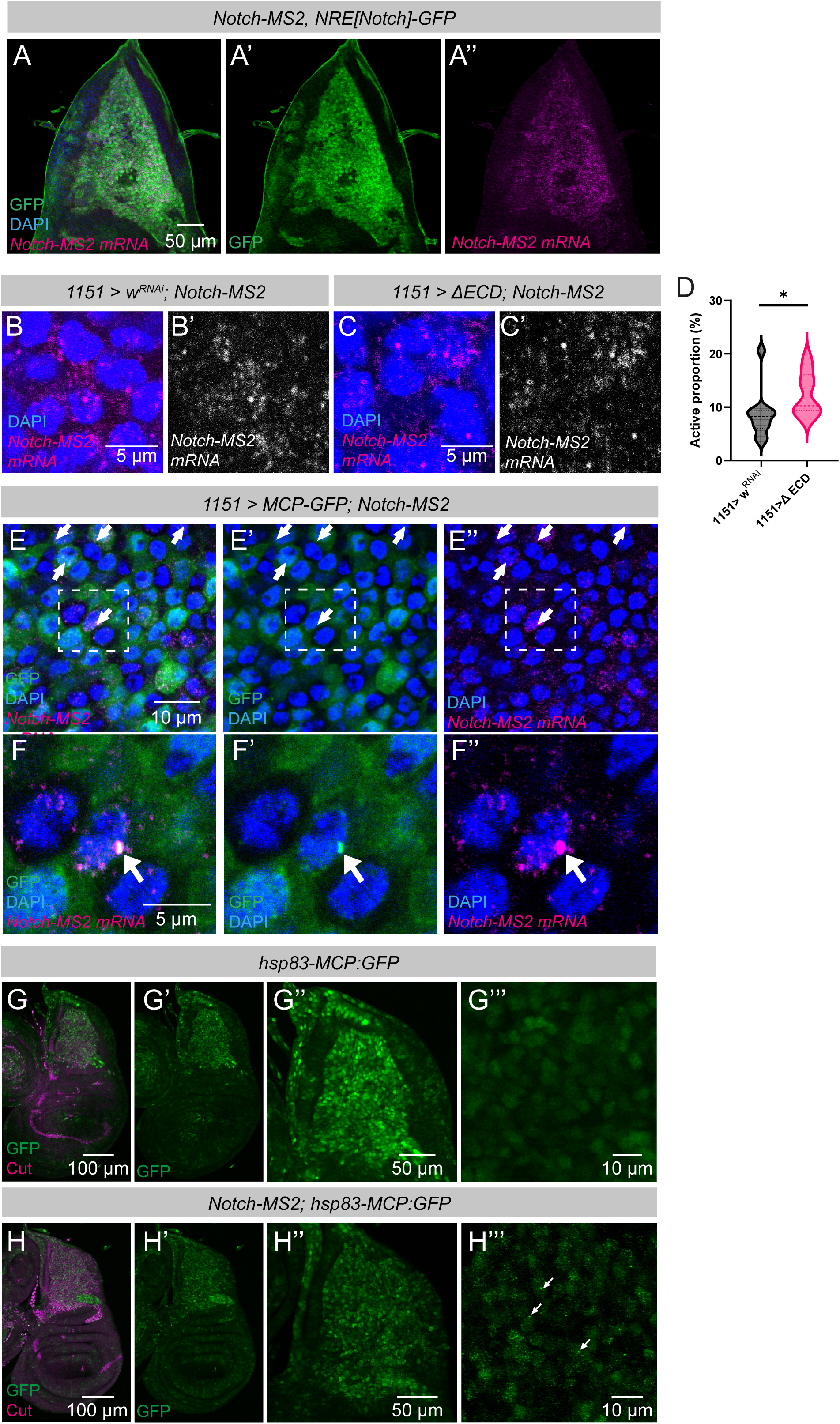
**(A-A’)** smFISH detection of Notch-MS2 transcripts using *yellow* probes (magenta) shows their colocalization with the Notch[NRE]-GFP reporter (green) in AMPs. **(B–D)** Detection of Notch-MS2 transcription by smFISH in AMPs under control (*1151 > w^RNAi^*) (A) and Notch-overactivation (*1151 > ΔECD*) (B) conditions. Nuclei are stained with DAPI (blue). Notch-MS2 transcripts are shown in magenta and grey. **(D)** Quantification of the proportion of transcriptionally active nuclei revealed a significant increase following Notch overactivation (*, *P* = 0.0159; Mann–Whitney test; control, *n* = 10 wing discs; ΔECD, *n* = 11 wing discs; three independent biological replicates). **(E–F″)** Detection of MCP-GFP (green) by immunostaining and Notch-MS2 transcription (magenta) by smFISH in AMPs. Nuclei are stained with DAPI (blue). **(F-F’’)** Higher-magnification images showing co-localization of MCP-GFP foci with active Notch-MS2 transcription sites within AMP nuclei. **(G–H‴)** Detection of Hsp83-MCP-GFP (green) and Cut (magenta) in AMPs in the absence (G–G‴) or presence (H–H‴) of the Notch-MS2 reporter. GFP accumulation is detected only in the presence of the Notch-MS2 reporter, confirming that MCP-GFP foci specifically mark active transcription sites.

## Notes

### Competing Interest Statement

The authors have declared no competing interest.

